# Enabling FIB-SEM Systems for Large Volume Connectomics and Cell Biology

**DOI:** 10.1101/852863

**Authors:** C. Shan Xu, Song Pang, Kenneth J. Hayworth, Harald F. Hess

## Abstract

Isotropic high-resolution imaging of large volumes provides unprecedented opportunities to advance connectomics and cell biology research. Conventional Focused Ion Beam Scanning Electron Microscopy (FIB-SEM) offers unique benefits such as high resolution (< 10 nm in x, y, and z), robust image alignment, and minimal artifacts for superior tracing of neurites. However, its prevailing deficiencies in imaging speed and duration cap the maximum possible image volume. We have developed technologies to overcome these limitations, thereby expanding the image volume of FIB-SEM by more than four orders of magnitude from 10^3^ µm^3^ to 3 x 10^7^ µm^3^ while maintaining an isotropic resolution of 8 x 8 x 8 nm^3^ voxels. These expanded volumes are now large enough to support connectomic studies, in which the superior z resolution enables automated tracing of fine neurites and reduces the time-consuming human proofreading effort. Moreover, by trading off imaging speed, the system can readily be operated at even higher resolutions achieving voxel sizes of 4 x 4 x 4 nm^3^, thereby generating ground truth of the smallest organelles for machine learning in connectomics and providing important insights into cell biology. Primarily limited by time, the maximum volume can be greatly extended.

Here we provide a detailed description of the enhanced FIB-SEM technology, which has transformed the conventional FIB-SEM from a laboratory tool that is unreliable for more than a few days to a robust imaging platform with long term reliability: capable of years of continuous imaging without defects in the final image stack. An in-depth description of the systematic approach to optimize operating parameters based on resolution requirements and electron dose boundary conditions is also explicitly disclosed. We further explore how this technology unleashes the full potential of FIB-SEM systems, revolutionizing volume electron microscopy (EM) imaging for biology by gaining access to large sample volumes with single-digit nanoscale isotropic resolution.

## Introduction

Connectomics aims to decipher the functions of brains by mapping their neural circuits. The findings will not only guide the next generation of development in deep learning and artificial intelligence but will also transform our understanding of the brain, in both healthy and diseased states. It can enhance research to find cures for brain disorders and perhaps even pave the way to ultimately comprehend the human mind.

Connectomics extracts the connectivity of neurons in the brain as a basis for understanding its function. The brain features that must be imaged with high resolution vary greatly in size, from the nm scale of the synapse to the mm length scale of the neurons that form even the smallest circuits. In three dimensions, these dimensional scales, taken to the third power, translate to 3D image stacks that can easily reach 10,000^3^ or Tera voxels. Furthermore, the data must maintain a high degree of accuracy and continuity, because even a few missing image planes could compromise the ability to reconstruct a neuron and trace its contribution to a neuronal network, invalidating months of data acquisition. These technical requirements have shaped the adaptation and development of FIB-SEM as a 3D volume imaging technique. The enhanced FIB-SEM technology developed at Howard Hughes Medical Institute’s Janelia Research Campus (***1***) has overcome the limitations of conventional volume electron microscopy methods, emerging as a novel approach for connectomic research.

FIB-SEM has been described previously in detail. It is a technique that has been used in materials science and the semiconductor industry for multiple decades. It has more recently been applied in biological imaging since 2006 (***2***). FIB-SEM uses scanning electron microscopy to raster scan the surface of a planar sample with a fine electron beam, a few nanometers in diameter, and monitors that surface by the back scattered electrons along with secondary electrons. Biological tissues are typically stained with heavy metals, such as osmium, which binds preferentially to the cell membranes and lipids thus enhancing the electron scattering signal at such locations. After imaging, the focused Ion beam (FIB), typically comprising 30 keV gallium ions, strafes across the imaged surface and ablates a few nanometers from the top of the sample to expose a new slightly deeper surface for subsequent imaging. Cycles of etching and imaging gradually erode away the sample while enabling the collection of a stack of consecutive 2D images, usually requiring tens of seconds to a few minutes per cycle.

Compared with serial thin section imaging (***3***, ***4***, ***5***), block-face based approaches (***1***, ***2***, ***6***, ***7***) provide greater consistency and stability of the image, thereby resulting in much better self-aligned image stacks. This is particularly important for connectomic studies: a well-registered and defect-free image stack is the foundation for successful automated segmentation. Serial sections incorporate folds and other imperfections in the sections thus posing significant challenges in registration and segmentation. Both diamond knife and FIB are options for the precise removal of material necessary for block-face based 3D imaging. FIB-based removal, unlike diamond knife, removes tissue at the atomic level without any mechanical moving components. It therefore offers the potential for nanometer control of the z-axis resolution. FIB milling is also less sensitive to damage by electron radiation from the SEM beam, so it can tolerate a higher electron dose to provide better signal-to-noise ratio (SNR) imaging (details of which are discussed in sections on Standard Resolution Mode and High Throughput Mode). The major disadvantages of conventional FIB-SEM include the slow imaging acquisition rate and lack of long-term stability. Additionally, the process of material ablation depletes the FIB gallium source in three to four days, so that just a few tens of microns in z thickness can be imaged before a pause is required to replenish the gallium source. After a pause, imprecise beam position could then result in excessive material loss while re-engaging the beam. Other factors such as room temperature fluctuations can disturb the fine control of the increment in z axis milling. These limitations constrain conventional FIB-SEM to small volumes.

To meet the large volume demands of connectomics, a number of enhancements to conventional FIB-SEM are discussed below. These enhancements have dramatically improved long-term reliability, and hence enabled uniform defect-free z increments that can support imaging of hundreds of microns in sample thickness. This represents a new regime in sample size and resolution for 3D volume imaging, with minimal trade-off between large volume and fine resolution. To probe biological questions with the most appropriate technology, it is necessary to characterize various platforms through a unified metric—minimum isotropic resolution, defined by the worst case in the x, y, or z axis. Because each imaging technology yields different resolutions in three axes, inferior resolution in any dimension can limit useful resolution in the other two, and impair the quality of subsequent image processing and analysis. Figure 1 provides an overview of the volume EM operating space. Specifically, it highlights the sample volume and minimum isotropic resolution that can be accessed only by long-term FIB-SEM imaging, and compares this to that by other volume EM imaging modalities. For example, diamond-knife cut serial section TEM with tomography, e.g. using ∼500-nm-thick sections, can give better spatial resolution at the cost of imaging volume, shown at the lower left of Figure 1. Conversely, even larger volumes with lower spatial resolution can be collected by diamond-knife cut serial section TEM or diamond-knife cut serial block-face SEM (SBFSEM), both enclosed for reference at the top of Figure 1. The space at the lower right of Figure 1 invites a technology for connectomic studies: the fine neurites can be traced at any random orientations without degradation of resolution, while neurons expanded over long distance can be fully captured by the large volume. Our enhanced FIB-SEM system delivers a larger volume with fine spatial resolution, addressing the need for connectomic studies. The red diagonal dotted lines indicate contours of constant imaging time: For a given time allotted for data acquisition, one may choose either finer resolution at the expense of sample volume or vice-versa. To explore this FIB-SEM application space, we break the regime into three domains, labeled simply: high resolution, standard resolution, and high throughput (reduced resolution). Application examples and opportunities of each will be discussed later.

**Figure 1.**
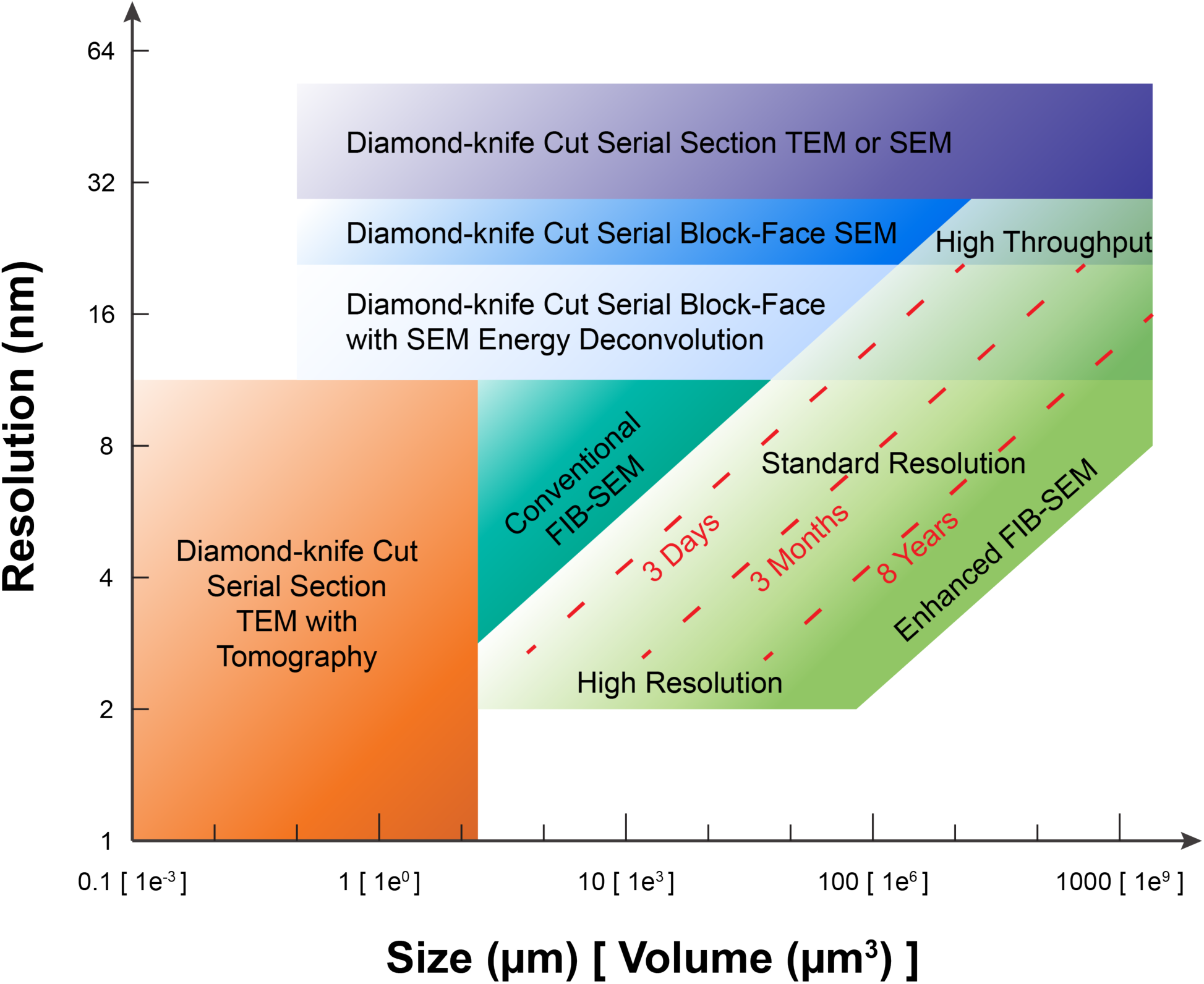
A comparison of different 3D volume EM imaging modalities in the application space defined by minimum isotropic resolution and total volume. The operating regime of enhanced FIB-SEM is divided into three zones: Standard Resolution, High Resolution, and High Throughput. The three red dotted lines indicate the general trade-off between resolution and total volume during FIB-SEM operations of 3 days, 3 months, and 8 years, respectively, using a single FIB-SEM system. These contours are sensitive to staining quality and contrast. The boundaries of the different imaging technologies outline the regimes where they have a preferential advantage, though in practice there is considerable overlap and only a fuzzy boundary.

The enhanced FIB-SEM system accelerates image acquisition while greatly improving reliability, advancing the operating period from days to years, and generating continuously imaged volumes larger than 10^7^ µm^3^ (***1***). These volumes are large enough for many connectomic studies, in which the excellent isotropic resolution enables automated tracing of small neurites and reduces the time-consuming human proofreading effort that is particularly crucial for dense reconstruction studies. The two most important technological advances are improvements in 1) imaging speed, and 2) system reliability, including error detection at all known failure modes and seamless recovery from those interruptions.

More than 10x improvement in imaging speed of the backscattered electron signal could be achieved without contrast degradation through a positive sample biasing strategy. In a Zeiss Gemini SEM column, this new configuration transforms the traditional in-column (InLens) detector into an effective backscattered electron detector (***1***). Compared with a traditional energy-selective backscattered (EsB) detector, the InLens detection via the biased scheme captures a larger fraction of the backscattered electrons, hence achieving a significant gain in imaging speed. Even higher throughput without sample bias is possible only when the steady state FIB-SEM imaging generates tolerable artifacts (e.g. artifacts like mild streaks which can be filtered out in the Fourier domain).

A major milestone in system reliability is accomplished through the following approaches: 1) using multiple layers of error and disturbance protection to prevent catastrophic failures; 2) comprehensive closed-loop control of the ion beam to maintain stability, allowing for a seamless restart of the imaging cycle after interruptions; 3) repositioning the FIB column to be 90 degrees from the SEM column to enable a shorter working distance and thus enhancing signal detection and image quality. Furthermore, features such as a zero overhead in-line image auto-optimization (focus, stigmation, and beam alignment) routinely ensure optimal images that are consistent throughout the entire volume. In depth descriptions of these improvements can be found in the Technology and Methods section of Xu et al. (***1***). Together, armed with innovative hardware architectures, system control designs and software algorithms, our enhanced system surpasses deficiencies in platform reliability against all known failure modes thereby overcoming the small volume limitations of conventional FIB-SEM techniques.

This transformative technology has empowered researchers to explore large volume connectomics and cell biology with an optimal balance of resolution, volume, and throughput.

The operating regimes can be grouped into three general categories. Details of seven exemplary cases including resolution requirement, FIB-SEM parameters, electron dose boundary, throughput estimate, and corresponding image reference are summarized in Table 1.

- Standard resolution (6–8 nm voxel) with and without parallel processing (by means of hot-knife partitioning), for large volume connectomics with traceability of small processes down to 15 nm; and for overview study of cellular structures.
- High resolution (< 5 nm voxel) for observing the finest details of synaptic ultrastructure in connectomic studies; and for fine structures of cell biology.
- High throughput at reduced resolution (> 10 nm voxel), for large volume connectomics with traceability of processes larger than 20 nm.

**Table 1.** Summary of FIB-SEM imaging conditions for Standard Resolution, High Resolution, and High Throughput Modes. Application examples and corresponding sample images are listed for reference. SEM electron landing energies used for electron energy dose estimate are 0.8 kV for High Resolution Mode and 1.2 kV for Standard and High Throughput Modes. SEM scanning rates are based on samples prepared by standard staining protocol. Faster scanning rates are expected samples prepared by improved staining protocol. (Throughput estimates include FIB milling and other operational overheads).

|  | Case 1 | Case 2 | Case 3 | Case 4 | Case 5 | Case 6 | Case 7 |
| --- | --- | --- | --- | --- | --- | --- | --- |
| Imaging mode | High Resolution | Standard Resolution | Standard Resolution | Standard Resolution | High Throughput | High Throughput | High Throughput |
| Application example | Mammalian single cell |  | *Drosophila* CNS |  |  |  | Mouse cortex |
| Embedding material | Epon with Durcupan cap |  |  | Durcupan |  |  |  |
| Hot-knife partitioning used | No | No | Yes | No | No | No | No |
| Voxel resolution (nm<sup>3</sup>) | 4 x 4 x 4 | 8 x 8 x 8 | 8 x 8 x 8 | 8 x 8 x 8 | 12 x 12 x 12 | 16 x 16 x 16 | 16 x 16 x 16 |
| SEM frame size (μm<sup>2</sup>) | 30 x 30 | 100 x 10 | 300 x 25 | 300 x 80 | 200 x 200 | 300 x 300 | 500 x 500 |
| SEM probe current (nA) | 0.25 | 2 | 3–4 | 3–4 | 4 | 4–6 | 8 |
| SEM scanning rate (MHz) | 0.2 | 0.25 | 2–3 | 2–3 | 4 | 4 | 8–12 |
| Electron dose (e<sup>-</sup>/voxel) | 7,800 | 50,000 | 6,200–12,400 | 6,200–12,400 | 6,200 | 6,200–9,300 | 4,100–6,200 |
| Electron radiation energy density (keV/nm<sup>3</sup>) | 97.7 | 117.2 | 14.6–29.3 | 14.6–29.3 | 4.3 | 1.8–2.7 | 1.2–1.8 |
| FIB milling probe (nA) | 15 | 15 | 15 | 15–30 | 15–30 | 30 | 30 |
| Throughput (μm<sup>3</sup>/month) | 0.03 x 10<sup>6</sup> | 0.2 x 10<sup>6</sup> | 1–1.5 x 10<sup>6</sup> | 1.8–2.4 x 10<sup>6</sup> | 12 x 10<sup>6</sup> | 24 x 10<sup>6</sup> | 33–50 x 10<sup>6</sup> |
| Sample image | Figure 5 | Figure 4 | Figure 9a | Figure 9a | Figure 9b | Figure 9c | Figure 9d |

## Methods

In this section, we present sample datasets from the *Drosophila* Central Nervous System (CNS), mammalian neural tissue, cultured mammalian cells, and the green algae *Chlamydomonas reinhardtii* to illustrate the power of this novel high-resolution and high throughput technique to address questions in both connectomics and cell biology. We explicitly report exemplary protocols in the three operating regimes with the goal of serving as a reference for readers to explore and optimize parameters for their own applications.

### Standard Resolution Mode

The initial motivation of our FIB-SEM platform development was mainly to satisfy the requirements of Janelia’s *Drosophila* connectomic research (***8***, ***9***), and to allow connectome mapping of small pieces of mouse and zebrafish nervous systems. Connectome studies comes with clearly defined resolution requirements—the finest neurites must be traceable by humans and should be reliably segmented by automated algorithms (***10***). For example, the very finest neural processes in *Drosophila* can be as little as to ∼15 nm (***11***), although such dimensions are only seen in short twigs attached to long-distance neurites of stouter caliber (***12***). In the case of mouse cortex, the finest long-distance axons can shrink to ∼50 nm, while dendritic spine necks can shrink to ∼40 nm (***13***). These fundamental biological dimensions determine the minimum isotropic resolution requirements for tracing neural circuits in each case.

To optimize the FIB-SEM operating conditions for each studied connectome, besides the resolution requirement, it is crucial to consider the effects of electron dose and electron radiation energy density. Electron dose, the number of electrons per voxel, is a determining factor for both SNR and imaging throughput. Electron radiation energy density, the product of electron dose and electron beam energy, represents the amount of irradiated energy per unit volume. It has a direct impact on milling rate or z removal consistency, which will become relevant later in the discussion. For reliable automated segmentation of *Drosophila* datasets, using 8nm isotropic voxels, an electron dose of 6,000–12,000 e^-^/voxel (3–4 nA electron beam, 2–3 MHz imaging rate) is required to achieve a sufficient SNR. We use a 1.2 kV electron beam to achieve the desired z resolution. Together this means that the electron radiation energy density is 15–30 keV/nm^3^ during steady state imaging. In comparison, the mammalian brain has larger neurites than that of *Drosophila*. For test datasets of well stained mouse cortex, 16 nm isotropic voxels and an electron dose as low as 4,000 e^-^/voxel (1.2 keV/nm^3^) appear sufficient for reliable automated segmentation. However more rigorous tests must be performed to determine the optimal voxel resolution and electron dose for mammalian connectomics using FIB-SEM. Applications using a voxel resolution larger than 10 nm will be further explored in the High Throughput Mode section.

Utilizing these Standard Resolution Mode parameters, we have acquired large (by typical FIB-SEM standards, ∼1 x 10^6^ µm^3^) connectomic datasets spanning parts of the *Drosophila* optic lobe (***14***, ***15***), mushroom body (***16***), and antennal lobe (***17***). However, with attempts to extend the imaged volume’s depth in the direction of the FIB beam beyond approximately 80 µm, thick-thin milling wave artifacts arose on the trailing edge of the block thus impairing the z resolution requirement. Interestingly, this milling wave phenomenon is dependent upon the electron radiation energy density. That is, lowering the total electron radiation energy density (keV/nm^3^) lessens the magnitude of milling wave artifacts which, in turn, allows the imaged volume to be made considerably longer in the direction of the FIB beam. Conceptually, one could multiply the SEM scanning rate on the heavily stained high contrast samples, thus expanding the imaged volume significantly. This observation suggests that the wave artifacts result, at least in part, from electron beam-induced modification of the plastic resin on the block surface.

Moreover, such electron beam-induced artifacts seem to be inherent to all block-face based imaging techniques, manifesting as thick-thin alternations in SBFSEM. Electron beam-induced modification and its effect on SBFSEM diamond-knife sectioning has been studied (***18***). The reported electron dose of SBFSEM to achieve consistent 25 nm sectioning is limited to 7.3 e^−^/nm^2^, which equates to an electron radiation energy density of 0.73 keV/nm^3^ using a 2.5 kV beam. Significantly higher electron doses cause SBFSEM sectioning to alternate between thick and thin slices, as if the surface layer had become a hardened crust. Recalling the above discussion, the value of electron radiation energy density of 0.73 keV/nm^3^ is actually 40 times lower than the value used in our *Drosophila* connectomics FIB-SEM datasets, implying that significant electron radiation-induced surface modification occurs during our standard FIB-SEM runs, while our FIB milling is less sensitive to electron radiation damage. These values are also generally consistent with the literature on radiation-induced chemical modification of polymers suggesting that a significant percentage of chemical bonds are modified at energy levels above 1 keV/nm^3^ (***19***).

The sample volume for connectome studies is typically set to encompass a particular circuit of interest. We were interested in imaging an entire central complex and mushroom body of *Drosophila*, which required an imaging volume of ∼250 x 250 x 250 µm^3^. Comparable research on mammalian brains would require considerably larger volumes (***13***). Such studies are clearly well beyond the 80 µm limit of Standard Resolution Mode FIB-SEM discussed here. To overcome this limitation, we developed an ultrastructurally smooth thick partitioning approach whereby heavy metal-stained, plastic-embedded samples could be subdivided into 20 µm thick slabs (***20***). These thick slabs are subsequently re-embedded and mounted so that their minimum dimension is oriented in the direction of the FIB beam thus avoiding any milling wave artifacts. Each thick slab is FIB-SEM imaged separately and the resulting volume datasets are stitched together computationally.

To be effective, the cut surfaces of the slabs must be smooth at the ultrastructural level and have only minimal material loss. Specifically, for connectomic research, all long-distance processes must remain traceable across sequential slabs. In our hands, traditional approaches using vibratome slicing or room-temperature microtomy failed to meet these requirements. Instead, we modified an existing hot-knife microtomy procedure (***21***) to use a heated, oil-lubricated diamond knife (***20***). These modifications allowed us to section both *Drosophila* and mammalian brain tissue at up to 25 µm thickness with an estimated material loss between consecutive slabs of ∼30 nm—sufficiently minimal to allow us to trace long-distance neurites in both fly and mammal (***20***).

In our largest study to date, we used this hot-knife approach to section an entire male *Drosophila* ventral nerve cord (VNC) into twenty-five slabs, each at 25 µm thick. This volume, 220 x 200 x 600 µm^3^, (∼2.6 x 10^7^ µm^3^) in total, was imaged in parallel across six FIB-SEM machines in about six months. In addition, a female *Drosophila* “hemi-brain” that spans the entire central complex, a unilateral mushroom body and optical lobe, was sectioned in a sagittal plane into 20-µm-thick consecutive slabs (Figure 2). Thirteen such slabs were imaged in two FIB-SEM machines. The fully segmented “hemi-brain”, 250 x 250 x 250 µm^3^, (∼1.6 x 10^7^ µm^3^) in volume, containing ∼25 x 10^3^ neurons with ∼60 x 10^6^ synaptic connections, is considered to be the largest connectome in the world in terms of the number of neurons and synapses being traced.

**Figure 2.**
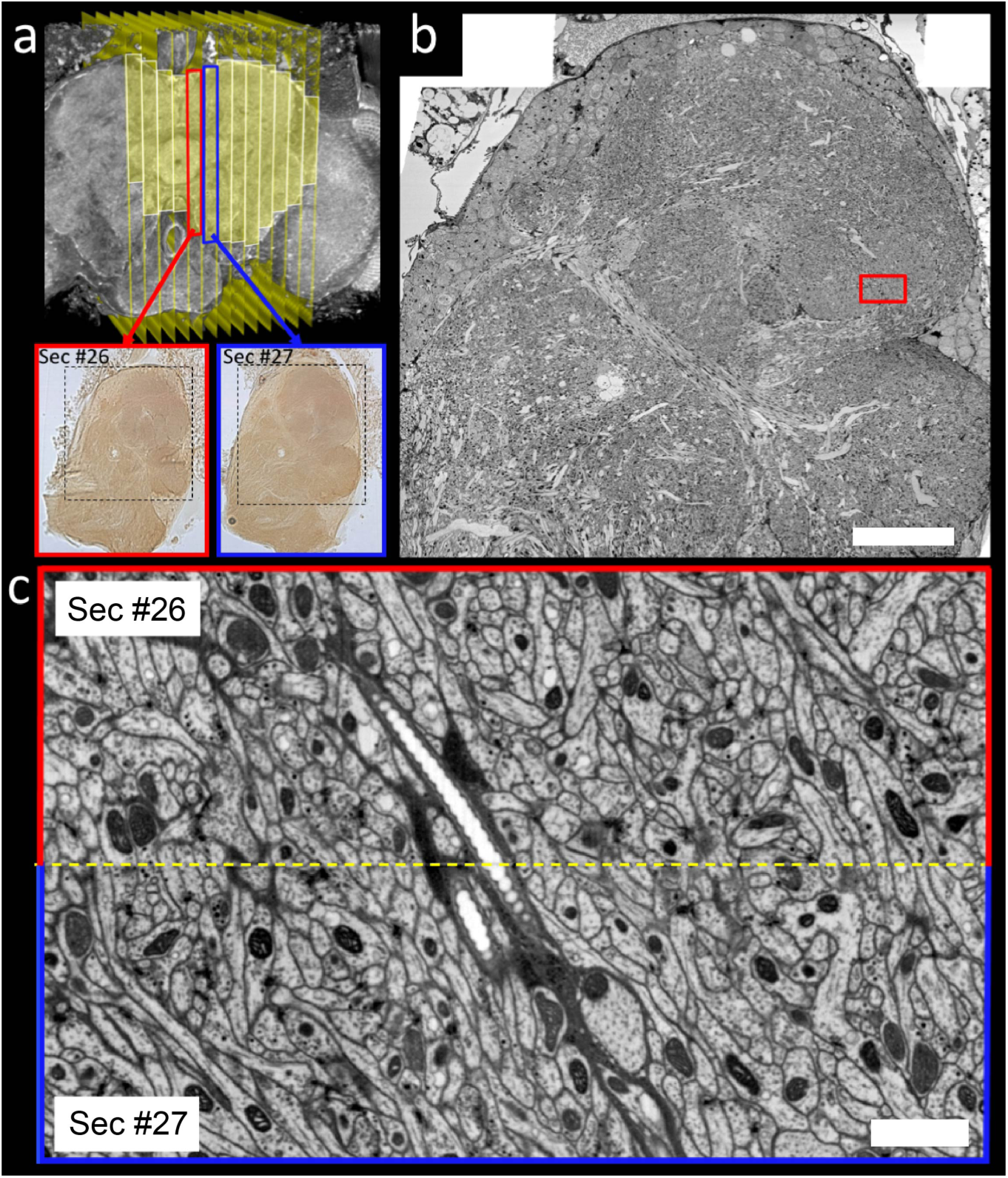
FIB-SEM imaging of an entire *Drosophila* central complex and a complete unilateral mushroom body. (a) X-ray micro-CT image of a *Drosophila* brain showing the locations of the thirteen consecutive 20 µm thick hot-knife sections that were FIB-SEM imaged for this study. Yellow highlighting is used to designate imaged volume. Light micrographs of two of these sections (labeled #26 and #27 in the overall series) are shown as well. Dashed boxes in the light micrographs designate regions that were FIB-SEM imaged. (b) Cross section through the FIB-SEM volume of Sec #26. Scale bar, 40 µm (c) Example zooming in on the boundary between hot-knife sections #26 and #27 whose FIB-SEM images have been computationally ‘volume stitched’. Yellow dashed line designates stitch line. The location of this stitched region is designated by the red rectangle in (b). Scale bar, 2 µm. Sample was prepared by Zhiyuan Lu (Dalhousie University)

The *Drosophila* connectome project demonstrates a significant advantage of the hot-knife approach—it allows many FIB-SEM machines to operate in parallel on a single imaging task. However, the limitations of this technique—that it is incompatible with heavy metal stained samples and Durcupan resin (***20***), unfortunately limits its adoption. As of today, hundreds of samples with smaller required volumes have been FIB-SEM imaged without hot-knife sectioning, while Durcupan, the preferred resin for FIB-SEM imaging, has been readily used for infiltration and embedding. Importantly, most recent improvements via the PLT-LTS progressive heavy metal enhancement staining protocol (***22***) enables even faster imaging rate without any degradation in quality (Figure 3). Lower radiation resulting from faster imaging significantly extends the FIB milling depth to hundreds of microns, thereby enabling much larger volume to be collected at standard resolution without the complexity of hot-knife partitioning. Thus, we have imaged an entire *Drosophila* L1 larval CNS (embedded in Durcupan) with 8 x 8 x 8 nm^3^ voxel resolution at 10 MHz, achieving synaptic-resolution without any gaps, all in less than 3 weeks. This dataset will be used to generate a full CNS connectome using automated methods. A typical *Drosophila* L1 larva CNS has a volume of ∼5 x 10^6^ µm^3^ in volume, and the faster SEM scanning rates enabled by the improved staining contrast allows us to extend the FIB milling depth to 200 µm with a sufficient margin. Primarily limited by time, the sample block in the z direction can be further expanded to its geometric limit. Armed with enlarged sample volumes and improved system throughput, we are embarking on a journey to tackle many biological questions. Promisingly, the 3D ultrastructure of tissue from the adult mouse hippocampus revealed by the enhanced FIB-SEM, has validated non-concentricity in myelinated axons (***23***), and discovered membrane-bound lipid-dense structures in neurons (***24***).

**Figure 3.**
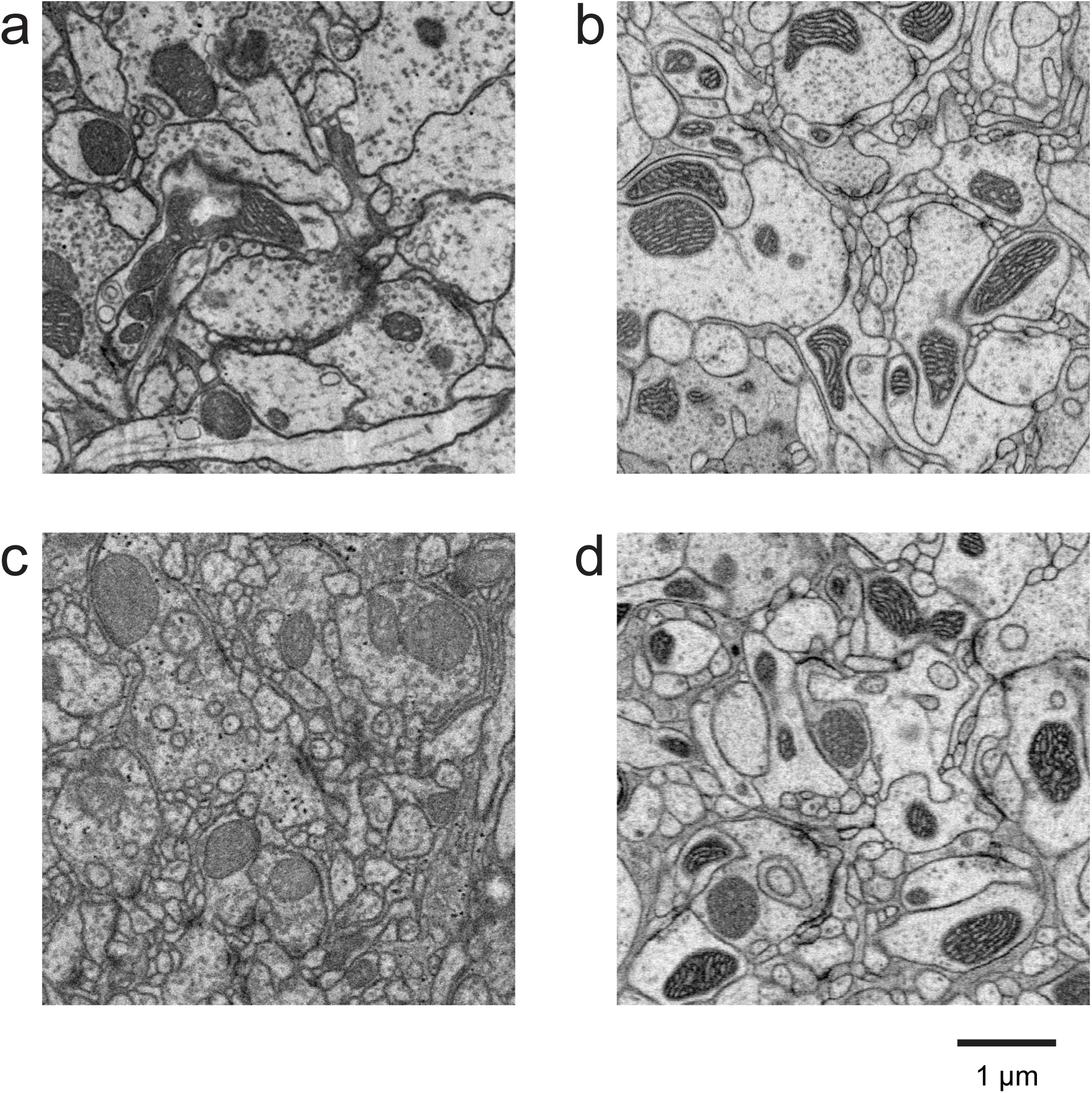
Faster imaging rate is achievable using improved staining *Drosophila* brain samples without image quality degradation. (a) standard staining sample 4 x 4 x 4 nm^3^ at 200 kHz. (b) improved staining sample 4 x 4 x 4 nm^3^ at 2 MHz. (c) standard staining sample 8 x 8 x 8 nm^3^ at 3 MHz. (d) improved staining sample 8 x 8 x 8 nm^3^ at 10 MHz. Samples were prepared by Zhiyuan Lu (Dalhousie University). PLT-LTS progressive heavy metal enhancement staining protocol (***22***) was used in improved staining samples.

In conjunction with connectomic studies, the improved imaging speed of our FIB-SEM system also allows rapid sampling of neuronal cultures or an entire mammalian cultured cell at 8 nm isotropic voxel in a week or less (***25***, ***26***). Note that the minimum dimension of cell samples grown on a cover glass is usually less than 20 µm, so that the imaging procedure can be simplified without hot-knife partitioning. Furthermore, a much higher electron dose of 50,000 e-/voxel can be used to boost SNR without uneven milling, the detailed condition of which is listed in Table 1 Case 2. This straightforward isotropic imaging mode opens up a new application space for cell biology, providing a better alternative to the serial section TEM or SEM cut on a diamond knife. The FIB-SEM datasets allow direct visualization of an entire cell in three dimensions, thus one no longer relies on sampling from 2D EM sections to infer its 3D organization. The comprehensive 3D overview of the structure and distribution of intracellular organelles permits examination of any arbitrary slices hence offering new insights, and could be mined for statistics. An exemplary dataset of a U2OS cell imaged by FIB-SEM at 8 x 8 x 8 nm^3^ voxels in 5 days is illustrated in Figure 4, where the nucleus, mitochondria, endoplasmic reticulum, and Golgi are visible in one slice plane cropped out of the 3D data volume. Furthermore, whole cell imaging at a standard resolution mode enables correlative light and electron microscopy (CLEM) applications, revealing a comprehensive picture of intracellular architecture, where subcellular components can be protein labeled and unknown EM morphologies can be classified without ambiguity (***26***). From the enlarged view in Figure 4b, one can see that the standard resolution is sufficient to resolve, for example, the cristae inside mitochondria. However, such resolution is challenged to distinguish, for example, between actin filaments and microtubules. In order to render finer details, we have therefore developed new capabilities for applications at a resolution finer than 6 nm.

**Figure 4.**
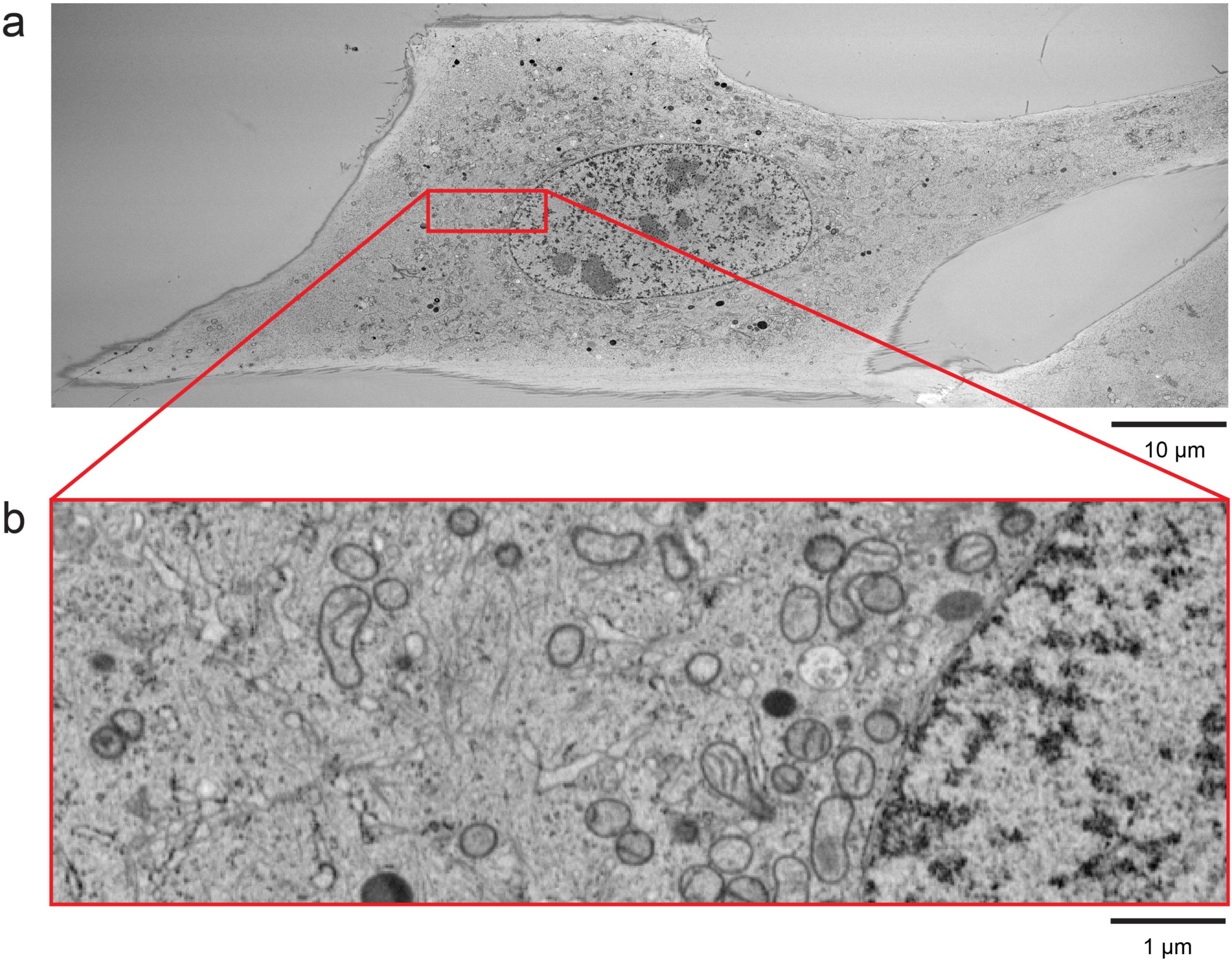
One slice plane from an isotropic image stack of a U2OS cell at 8 x 8 x 8 nm^3^ voxel resolution. (a) The overview of the entire cell shows various intracellular organelles and their distributions. (b) 10x zoom of the red box in (a) provides a closer view of mitochondria, Golgi, endoplasmic reticulum, and actin filaments near the nucleus. Scale bar, 10 µm in (a) and 1 µm in (b). Sample was prepared by Kathy Schaefer, David Hoffman, Gleb Shtengel, and Amalia H. Pasolli (Howard Hughes Medical Institute, Janelia Research Campus).

### High Resolution Mode

The requirement for a High Resolution Mode is considerably more time consuming per unit volume than that of the Standard Resolution Mode described above, while further improvements in resolution should enable important scientific advances by rendering finer details of cell biology and ground truth of connectomics. Even higher isotropic resolution is possible at the expense of imaging speed (or total volume within a given time), the significant challenges stem from the requirement to improve resolution in x, y, and z directions simultaneously.

To image the block-face of a sample with SEM, a sharply focused electron beam is raster scanned across the surface. Lateral resolution in the x y direction is dependent primarily on the blur of the incoming beam, and to a lesser extent on the lateral scattering that generates secondary or backscattered electrons (***27***). Additionally, the major contributions to the beam blur include spherical aberrations and Coulomb repulsion from the electron lens. The easiest way to mitigate this blur for finer x y resolution is simply to lower the beam aperture, with concurrent loss of imaging current and thereby imaging speed. The z resolution, on the other hand, is dependent upon the incoming electron landing energy that determines its probing depth (***1***). For a typical osmium-stained resin-embedded biological sample, a landing energy of 800 to 1200 eV offers a z-axis resolution of ∼5 to 8 nm at contrasts of 20% to 40%, respectively. Contrast between the heavy metal stain and the background signal of embedding resin deteriorates rapidly if energies below 800 eV are used. Even higher z resolution can be achieved by lowering electron landing energy to reduce the point spread function size along z-axis at the cost of reduced contrast.

Here we present typical operating conditions for the High Resolution Mode. Lateral resolutions of 1.5–3 nm (using the 25%–75% edge transition definition) should be possible with commercial SEMs operating at beam currents of 0.2–0.3 nA. A corresponding sampling interval of 2–4 nm is then used to match such high spatial detail. Similar to the SNR requirement in Standard Resolution Mode, an electron dose of approximately 6,000–8,000 e^-^/voxel is needed to achieve reasonable contrast in the High Resolution Mode, which inevitably relies on the heavy metal staining level. With an average staining contrast, using a 0.8 kV 0.25-nA electron beam, such a dose corresponds to a scanning rate of ∼200 kHz, which translates to a 30 x 30 x 30 µm^3^ (∼2.7 x 10^4^ µm^3^) volume in a month at 4 x 4 x 4 nm^3^ voxels. Note that the electron radiation energy density of this imaging condition at ∼97.7 keV/nm^3^ is more than triple compared with that of the Standard Resolution Mode (Case 1 vs. Cases 3 & 4 in Table 1). As a result, the imaged volume depth in the direction of FIB beam without milling artifacts drops to ∼20–30 µm, but this is manageable since the minimum dimension of cultured cells (typically less than 20 µm) can be aligned to the FIB milling direction. Ultimately, this mode opens up a unique application space that compliments standard EM tomography which can achieve higher spatial resolution at the cost of smaller and less thick samples. Although the tomographic approach can be extended to thicker effective samples by stitching multiple samples together, it takes considerable effort with diminishing returns for stitching a larger number of sections, compared with the ease of the FIB-SEM approach.

Figure 5 shows a typical image of a portion of a HeLa cell that was grown on a cover glass, then high pressure frozen with standard freeze substitution, osmium tetroxide staining, and Durcupan resin embedding. After being trimmed to a pedestal of ∼80 x 80 x 80 µm^3^ in size, it was FIB-SEM imaged over a 50 x 8 µm^2^ region (8 µm dimension of imaging aligned with the direction of the FIB beam) with 4 x 4 x 4 nm^3^ voxels at 200 kHz, using an electron landing energy of 1 keV. The Figure exemplifies the resolution and quality that can be achieved with this technique. The nuclear envelope in both a perpendicular and a tangential slice plane shows the double membrane and nuclear pores. Chromatin is also visible as the dark granular structure inside the nucleus. On the surface of the tangential plane, nuclear membrane poly-ribosome chains are visible. The resolution is sufficient to see the hollow center of the 20 nm diameter microtubules that lay on the surface of the nuclear membrane. They are easily identified as parallel lines when bisected by the image plane along their axis. The same is true of the microtubule structure of the centrosome. Golgi, endoplasmic reticulum, mitochondria, etc. are all identifiable in the cytoplasm. Clearly, these structures are much better resolved than those using a standard resolution of 8 x 8 x 8 nm^3^ shown in Figure 4. Additional examples, such as neural tissues and single cells of higher resolution FIB-SEM datasets are presented in the references (***1***, ***16***, ***28***). Figure 6 offers a side-by-side comparison of the High Resolution Mode over the Standard Resolution Mode. Figure 7 demonstrates the power of high resolution datasets in revealing and classifying 3D cellular structures, and such details could be objectively quantified and extracted for statistical purposes. Likewise, these high resolution images can also help to detail typical synaptic morphology and aid in deciphering the extremely fine processes of neural connectivity (Figure 8) in connectomic studies.

**Figure 5.**
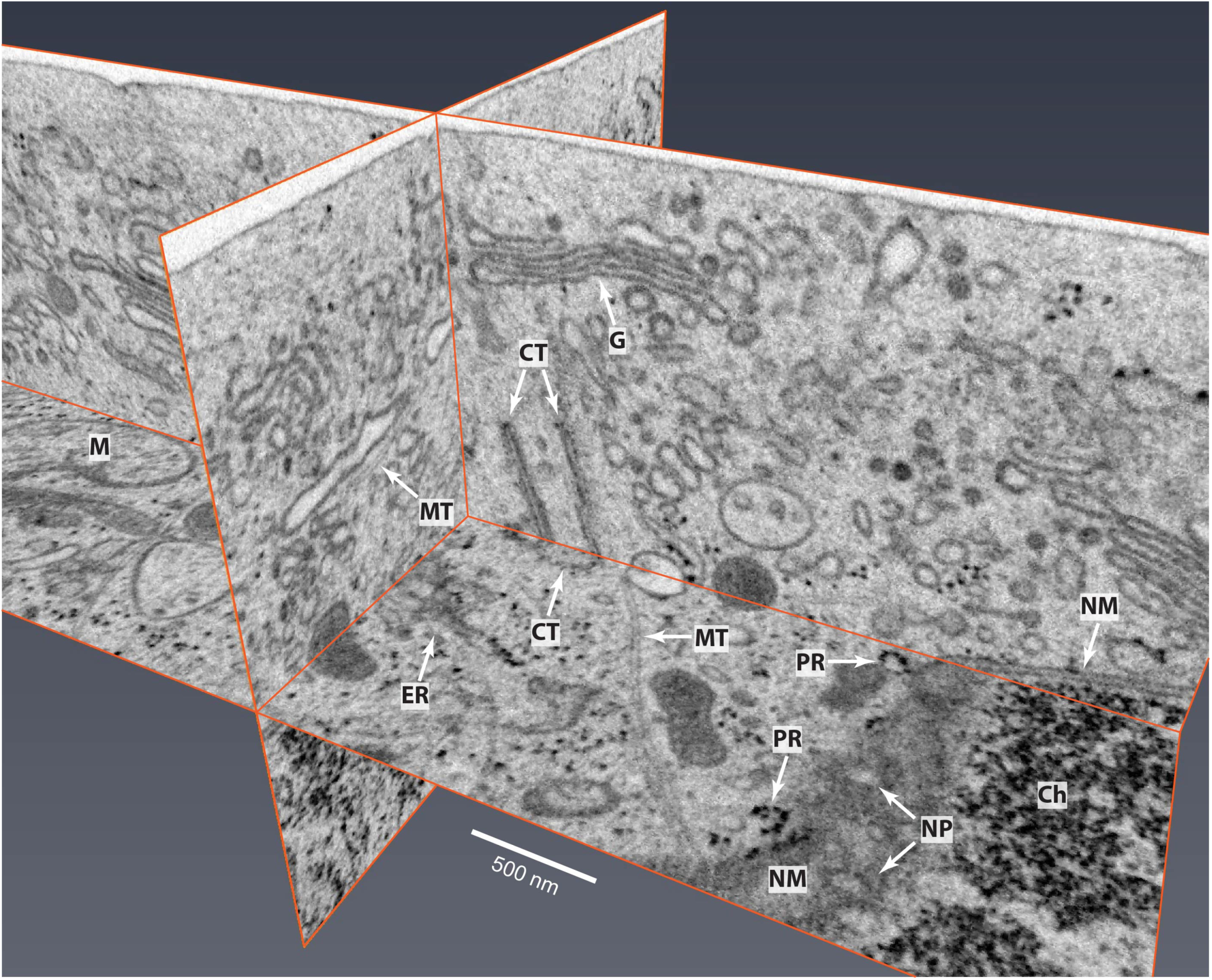
FIB-SEM image of a HeLa cell using High Resolution Mode at 1 keV, 0.25 nA, 200 kHz, and 4 x 4 x 4 nm^3^ voxels. Many intracellular cellular organelles are clearly resolved. They are labeled as: chromatin (Ch), centrosome (CT), endoplasmic reticulum (ER), Golgi (G), mitochondrion (M), microtubule (MT), nuclear membrane (NM), nuclear pore (NP), and polyribosome (PR). Sample was prepared by Aubrey Weigel, Gleb Shtengel, and Amalia H. Pasolli (Howard Hughes Medical Institute, Janelia Research Campus).

**Figure 6.**
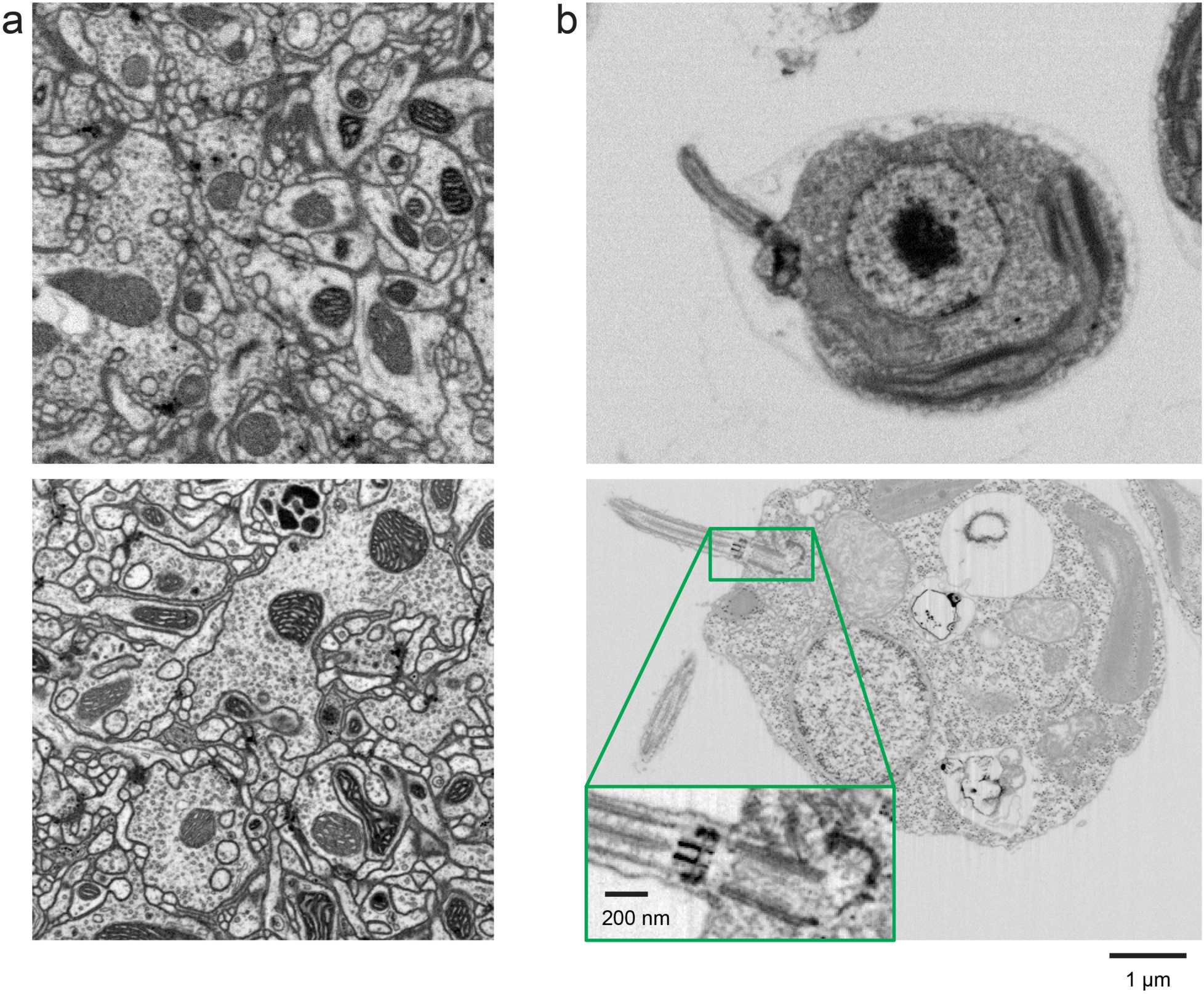
Improved FIB-SEM resolution reveals more detailed cellular structures in biological samples. Typical images of (a) *Drosophila* central complex and (b) *Chlamydomonas reinhardtii*, using standard 8 x 8 x 8 nm^3^ voxel imaging condition are shown in the top panels. The bottom panels show the corresponding high-resolution images at 4 x 4 x 4 nm^3^ voxels. Scale bar, 1 µm. Inset scale bar, 200 nm. Reproduced from (***1***)

**Figure 7.**
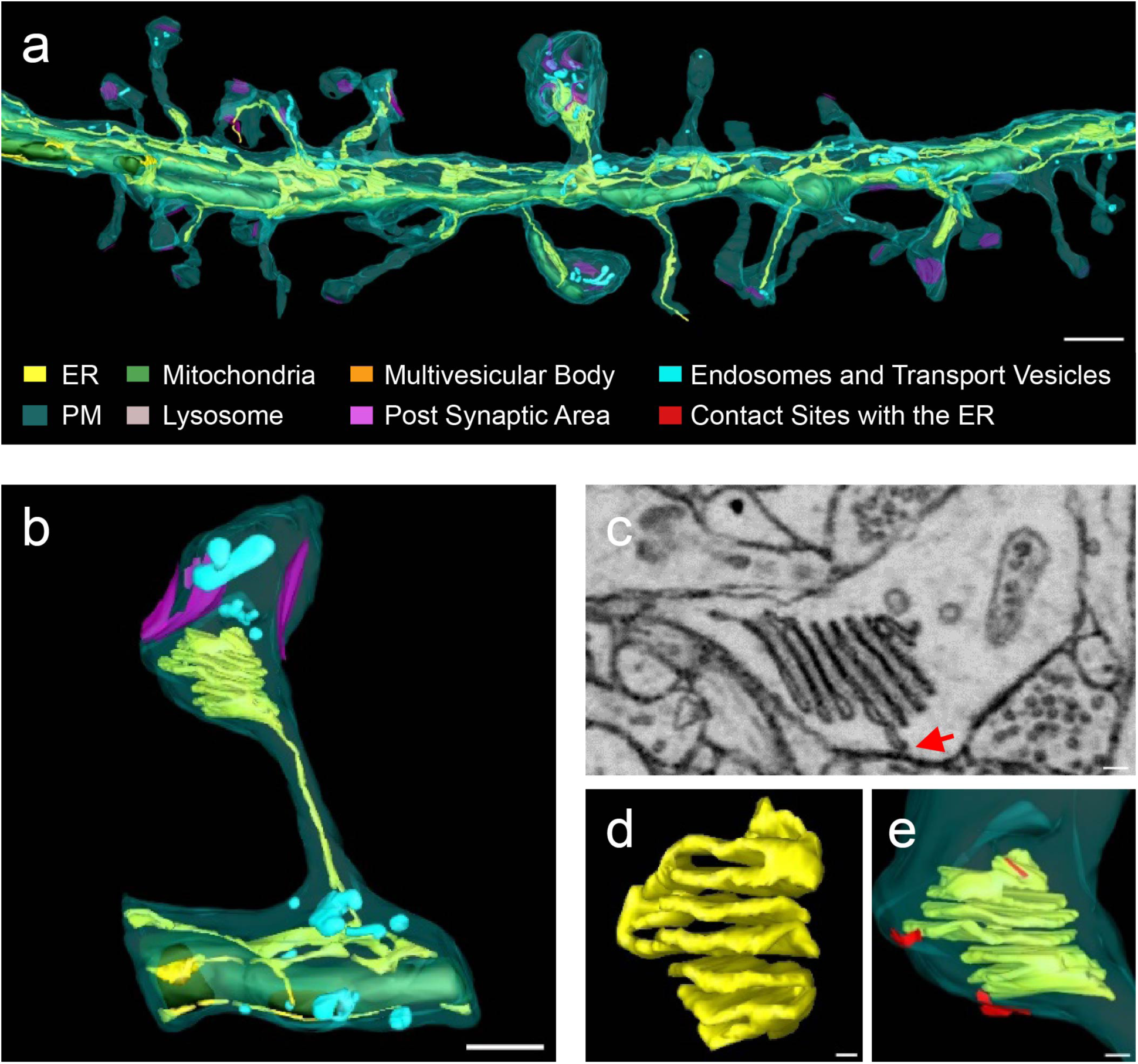
Visualization of 3D structures of a dendritic segment using high resolution 4 x 4 x 4 nm^3^ resolution FIB-SEM data. (a) 3D model showing all membranous organelles present in a dendrite: endoplasmic reticulum (ER), plasma membrane (PM), mitochondria, lysosome, multivesicular body, postsynaptic area, endosomes and transport vesicles, and contract sides of PM with ER are highlighted in red. (b) Zoomed in view of one dendritic spine. (c) Single FIB-SEM image showing a cross-section of the spine apparatus, which comprises seven cisternae, one of which makes a contact with the PM in the plane of the image (red arrow). (d) 3D reconstruction of the spine apparatus shown in (c). (e) Two contacts of spine apparatus with the PM (red). Scale bar, 800 nm in (a), 400 nm in (b), and 80 nm in (c)–(e). Reproduced from (***28***)

**Figure 8.**
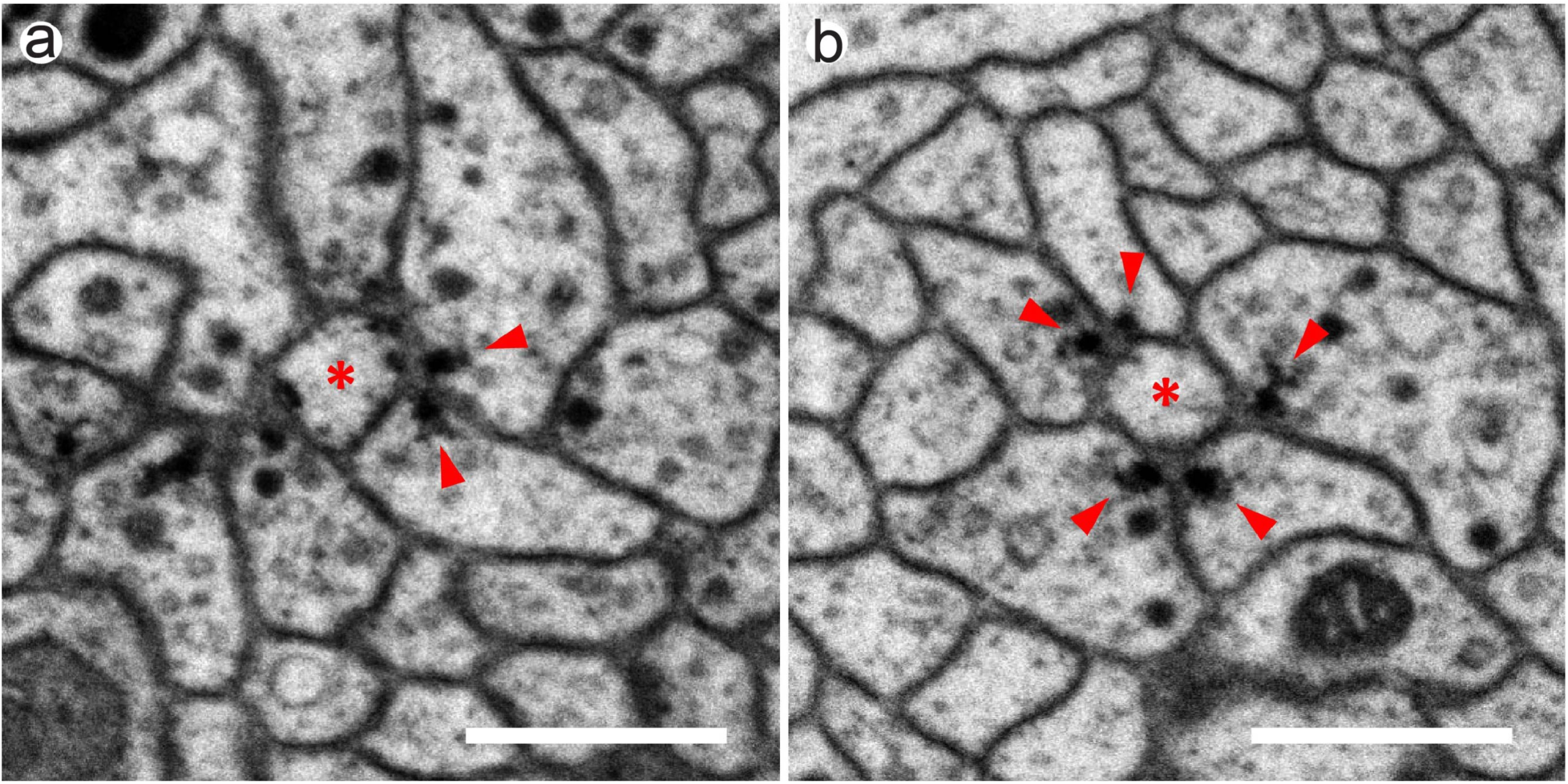
Images from the higher resolution dataset allow catalog of various newly observed synaptic motifs in *Drosophila* mushroom body with greater confidence. (a) A triangular motif of two adjacent Kenyon cells synapse onto a mushroom body output neuron. (b) A rosette motif of five Kenyon cells surround a mushroom body output neuron. Kenyon cells and mushroom body output neurons are labeled by red arrowheads and red asterisk, respectively. Scale bars, 500 nm. Reproduced from (***16***)

With the recently improved staining protocol (***22***), it is encouraging that without imaging degradation we could demonstrate an additional 10x improvement in the SEM scan rate: from 200 kHz (Figure 3a) to 2 MHz (Figure 3b) at 4 x 4 x 4 nm^3^ using a *Drosophila* brain sample. At such a rate, we have imaged sub-compartments of the *Drosophila* brain rather rapidly. For example, a fan-shaped body middle column, 40 x 50 x 50 µm^3^ (∼1 x 10^5^ µm^3^), was imaged in two weeks, while the mushroom body, 50 x 50 x 120 µm^3^ (∼3 x 10^5^ µm^3^), could be potentially imaged within a month, all at 4 x 4 x 4 nm^3^ resolution. Evidently, a successful strong staining protocol can significantly benefit high-resolution image acquisition because the faster imaging rate substantially improves the throughput.

In summary, high-resolution datasets, while limited in volume, can provide additional important scientific details, and serve as an accurate gold standard to aid in the interpretation of much larger volume datasets obtained through Standard Resolution and High Throughput Modes. Furthermore, such datasets can potentially serve as ground truth for machine learning.

### High Throughput Mode

With an initial focus on the *Drosophila* connectome, we have chosen 8 nm isotropic voxel datasets as our standard operating mode to enable tracing *Drosophila*’s very fine neurites. To meet the demands of diverse collaborations, we have expanded 3D FIB-SEM operation space to High Throughput Mode, in which a larger volume can be FIB-SEM imaged at a 10x or more improvement in volume rate with sufficient resolution to yield biologically significant details for select questions.

FIB-SEM ability to acquire an isotropic 3D dataset enables visualization of uniform re-sliced planes at any random angle and permits extension to resolutions of any voxel size. One can choose larger (> 10 nm) voxels, in which the imaging rate is substantially accelerated by the cubic power reduction in the number of voxels. Additionally, higher SEM probe currents can be used for faster scan rates without sacrificing SNR. In theory, either larger voxels or faster imaging with lower electron dose and SNR can translate into higher volume imaging rates. To determine the boundary conditions, neuron traceability as a function of voxel size and SNR needs to be carefully evaluated. We have generated a dataset of the *Drosophila* brain with various voxel resolutions from 8 x 8 x 8 nm^3^ to 16 x 16 x 16 nm^3^ and electron doses from ∼3,000 e^-^/voxel to ∼12,000 e^-^/voxel. Examples and corresponding images of selected conditions are shown in Table 1 and Figure 9, respectively. A comparison is drawn among Standard Resolution Mode with (Case 3) and without (Case 4) hot-knife processing, and High Throughput Mode without hot-knife (Cases 5 & 6). We have imaged multiple *Drosophila* larval CNS (L1 and L3) samples (embedded in Durcupan) at 12 x 12 x 12 nm^3^ voxel resolution for tracing the skeletons of all neurons. The typically volumes of L1 or L3 are about 5 x 10^6^ µm^3^ or 1 x 10^7^ µm^3^, respectively. For samples prepared by our standard staining protocol, at an SEM scanning rate of 3 MHz, it took roughly 20 days for L1, and 40 days for L3. For the recent improved-staining samples, at an SEM scanning rate of 10 MHz, we finished several L1 samples in 10 days each, and expect an L3 sample in 20 days.

**Figure 9.**
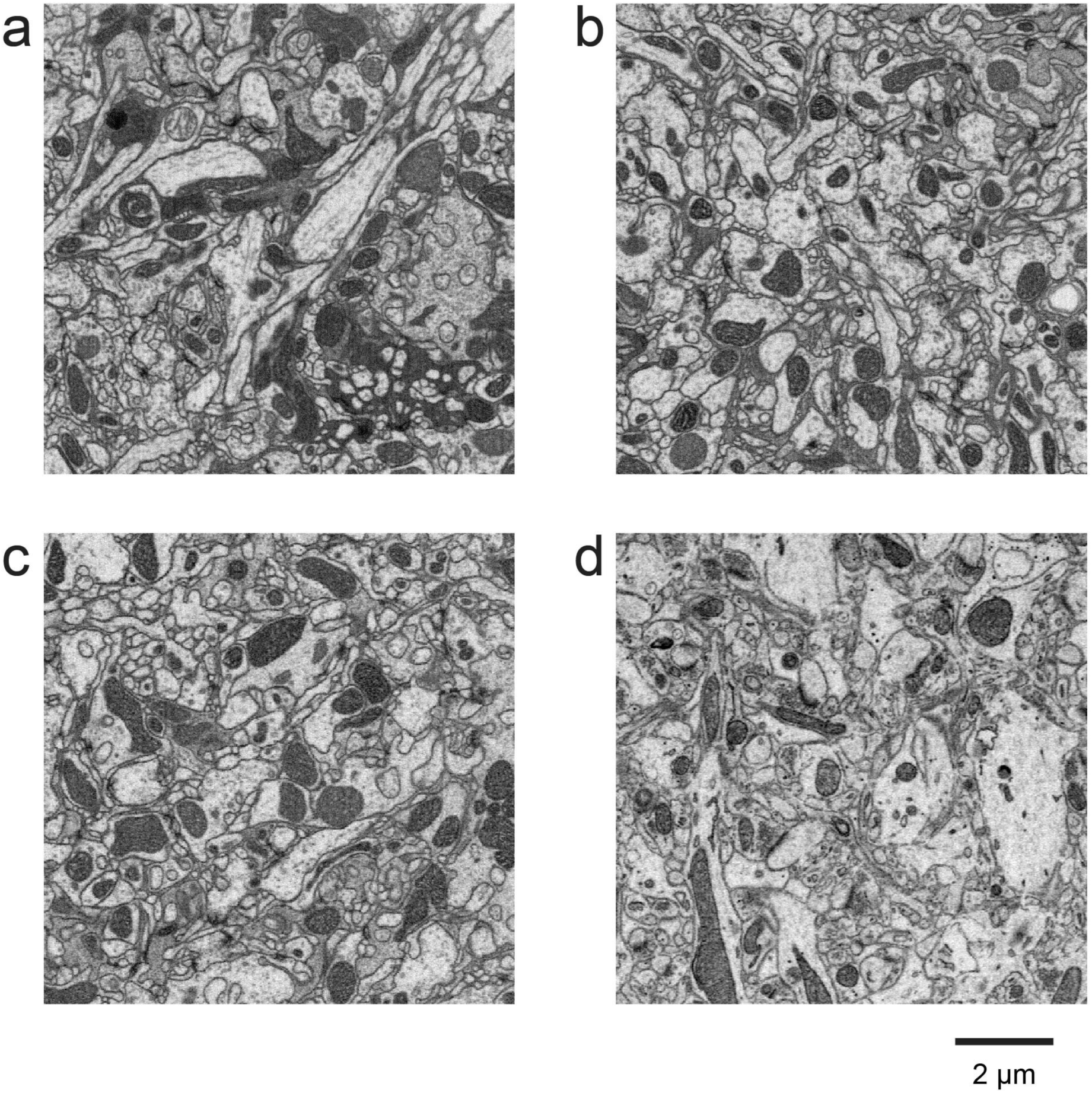
Sample images of connectomic studies using the enhanced FIB-SEM system. SEM probe landing energy was fixed at 1.2 kV. (a) *Drosophila* brain with 8 x 8 x 8 nm^3^ voxels scanned by a 4-nA SEM probe at 3 MHz. (b) *Drosophila* brain with 12 x 12 x 12 nm^3^ voxels scanned by a 4-nA SEM probe at 4 MHz. (c) *Drosophila* brain with 16 x 16 x 16 nm^3^ voxels scanned by a 4-nA SEM probe at 4 MHz. (d) Mouse cortex with 16 x 16 x 16 nm^3^ voxels scanned by an 8-nA SEM probe at 12 MHz. Scale bar, 2 µm. *Drosophila* brain samples were prepared by Zhiyuan Lu (Dalhousie University) and mouse cortex sample was prepared by Graham Knott (École Polytechnique Fédérale de Lausanne).

Even though *Drosophila* brain tissues are usually more challenging to image because of their lower staining contrast and smaller processes compared with mammalian neural tissues, it is encouraging that not only neurites as small as 50 nm in diameter are well distinguished and traceable, but also T-bar synapses are clearly visible using High Throughput Mode in all test conditions. The readily extractable neuron shapes can also aid in cell type identification in comparison with those obtained from optical images. Moreover, analysis of automated segmentation and human proofreading suggests that neuron tracing is more sensitive to isotropic voxel size than SNR. Further increases in voxel size and SEM scanning rate have diminishing returns for volume throughput once the volume rate of SEM imaging exceeds that of FIB milling. For case 7 in Table 1 as an example, with a voxel resolution of 16 x 16 x 16 nm^3^, the SEM volume image rate is close to the maximum FIB removal rate of ∼50 µm^3^/s using a 30-nA milling probe. Therefore, while all conditions yield traceable results, we recommend not to exceed a 16 x 16 x 16 nm^3^ voxel size for connectomic studies prepared for automated segmentation. Ultimately, it is a delicate balance between traceable resolution and throughput.

Once the resolution boundary conditions of High Throughput Mode have been determined, it is imperative to characterize the corresponding electron dose and radiation energy density. As shown in Table 1, the electron doses used in the three operating modes (for all cases except Case 2) remain rather consistent at ∼6,000 e^-^/voxel due to the SNR requirement. In order to accommodate large volumes, the electron radiation energy density decreases monotonically from 97.7 keV/nm^3^ to 1.2 keV/nm^3^ as the voxel size increases from 4 nm to 16 nm. We discover that the reduction of electron radiation damage to specimen widens the margin of uniform milling significantly, which in turn allows larger dimension samples to be directly imaged without the complication and overhead from hot-knife partitioning and post data stitching. Consequently, the material loss at each hot-knife interface (∼30 nm) can be completely avoided. For example, to image the entire *Drosophila* VNC, a volume of 220 x 200 x 600 µm^3^, (∼2.6 x 10^7^ µm^3^), requires roughly two FIB-SEM-years at 8 x 8 x 8 nm^3^ voxel resolution in Standard Resolution Mode with the hot-knife procedure; in contrast, it can be accomplished efficiently without hot-knife using High Throughput Mode (Cases 5 & 6 in Table 1), primarily for tracing the lower order branches of all neurons. Moreover, the faster scanning rate enabled by the recently improved staining protocol extends the FIB milling depth beyond 200 µm, therefore such a volume can be accomplished in two FIB-SEM-months at 12 x 12 x 12 nm^3^, or less than five FIB-SEM-months at 8 x 8 x 8 nm^3^ voxel resolution with minimal milling artifacts.

Importantly, while High Throughput Mode overlaps with SBFSEM in the resolution-volume space (Figure 1), FIB-SEM provides higher SNR images than SBFSEM, because FIB milling is less sensitive to electron radiation damage compared with the diamond-knife cutting. To obtain consistent cutting, SBFSEM caps the electron dose and radiation energy density at 2,000 e^-^/voxel and 0.73 keV/nm^3^, respectively (***18***). The 3x or more electron dose in FIB-SEM directly translates into higher SNR images thereby enabling higher accuracy in the subsequent analysis and interpretation.

Altogether, the High Throughput Mode offers biologists a viable option to obtain an overview of large volume neural circuitry in a reasonable amount of time. This regime should be particularly attractive to researchers who focus on structures larger than 16 nm and who are interested in mining statistical information from multiple samples. The speed improvement of High Throughput Mode is rather promising: the same volume of tissue can be imaged at least 10x faster than that of Standard Resolution Mode. Coupled with the staining improvement, the High Throughput Mode opens up exciting new avenues for connectomic studies. As can be seen in Figure 9d, thanks to a much improved staining contrast of mammalian brain tissues (***29***), we are able to raise the SEM imaging rate to 12 MHz using an 8-nA electron beam without image degradation (Table 1, Case 7). Under such conditions, the projected imaging time of a 1 mm^3^ volume is only 2–3 FIB-SEM-years, a significant improvement over nearly one FIB-SEM-century using the Standard Resolution Mode!

## Summary

Three-dimensional imaging offers tremendous value to elucidate biological structures and decipher their function. The enhanced FIB-SEM technology has addressed the limitations of existing imaging modalities thus effectively expanding the operating space of volume EM, delivering fine isotropic resolution, and high throughput with long-term reliability to image sufficiently large volumes encompassing the entire region of interest. The expanded volumes open a vast new regime in scientific learning, where nano-scale resolution coupled with meso and even macro scale volumes is critical. The largest connectome in the world has been generated using this enhanced FIB-SEM platform, where the superior z resolution empowers automated tracing of neurites and reduces the time-consuming human proofreading effort. Increased resolution further improves the interpretation of otherwise ambiguous details. Nearly all organelles can be resolved and classified with whole cell imaging at 4 nm voxel resolution. We have routinely imaged entire mammalian cells at this resolution to study the close contacts among various organelles. Furthermore, new CLEM applications enabled at the whole-cell level, can readily probe cell biology questions that are otherwise intractable.

At the forefront of volume EM imaging innovations, enhanced FIB-SEM technology pushes the envelope of image acquisition capability and system reliability, offering a novel package suited for large volume connectomics and cell biology.

## Acknowledgements

We would like to thank David Peale and Patrick Lee for consulting support in system modification. We also thank Zhiyuan Lu, Gleb Shtengel, David Hoffman, Amalia H. Pasolli, Kathy Schaefer, Aubrey Weigel, Nadine Randel, and Michael J. Winding for EM sample preparation. We gratefully acknowledge Patrick Naulleau, Ian A. Meinertzhagen, and Steve Plaza for reviewing the manuscript and providing timely feedback. Our gratitude extends to Janelia FlyEM connectome program, in particular Gerry Rubin and Steve Plaza for their leadership. We were solely funded by the Howard Hughes Medical Institute.

